# Benchmarking of Small Molecule Feature Representations for hERG, Nav1.5, and Cav1.2 Cardiotoxicity Prediction

**DOI:** 10.1101/2023.08.15.553429

**Authors:** Issar Arab, Kristof Egghe, Kris Laukens, Ke Chen, Khaled Barakat, Wout Bittremieux

## Abstract

In the field of drug discovery, there is a substantial challenge in seeking out chemical structures that possess desirable pharmacological, toxicological, and pharmacokinetic properties. Complications arise when drugs interfere with the functioning of cardiac ion channels, leading to serious cardiovascular consequences. The discontinuation and removal of numerous approved drugs from the market or at late development stages in the pipeline due to such inhibitory effects further highlight the urgency of addressing this issue. Consequently, the early prediction of potential blockers targeting cardiac ion channels during the drug discovery process is of paramount importance. This study introduces a deep learning framework that computationally determines the cardiotoxicity associated with the voltagegated potassium channel (hERG), the voltage-gated calcium channel (Cav1.2), and the voltage-gated sodium channel (Nav1.5) for drug candidates. The predictive capabilities of three feature representations—molecular fingerprints, descriptors, and graph-based numerical representations— are rigorously benchmarked. Additionally, a novel training and evaluation dataset framework is presented, enabling predictive model training of drug off-target cardiotoxicity using a comprehensive and large curated dataset covering these three cardiac ion channels. To facilitate these predictions, a robust and comprehensive small molecule cardiotoxicity prediction tool named CToxPred has been developed. It is made available as open source under the permissive MIT license at https://github.com/issararab/CToxPred.

## 1. Introduction

Drug discovery is a complex and multifaceted process that involves the identification and development of new therapeutic agents to treat various diseases. It encompasses a range of scientific disciplines, including chemistry, biology, pharmacology, and medicine, and it involves the discovery, design, synthesis, and optimization of new chemical entities that can modulate disease targets in a safe and effective manner. Conventionally, within the realm of drug discovery, researchers undertake *in vitro* and *in vivo* studies to assess the pharmacodynamics and pharmacokinetic (PD/PK) characteristics of chosen candidates derived from preliminary screening outcomes [1][2]. These experiments are not only demanding in terms of time and financial resources but also raise ethical concerns, particularly in cases involving animal testing [3]. Previous research has demonstrated that the creation of new drugs is a time-consuming and elaborate process that on average can take from six to twelve years and involve expenses of up to 2.5 billion dollars [4][5][6]. Out of this substantial financial cost, approximately 1.1 billion dollars is allocated for the stages of drug development preceding human trials [6].

Among the five essential pharmacokinetic attributes—chemical absorption, distribution, metabolism, excretion, and toxicity—is toxicity, which necessitates strict validation prior to granting clinical trial approval for a novel drug candidate [7]. According to the directives outlined by the International Conference on Harmonization of Technical Requirements for the Registration of Pharmaceuticals for Human Use, there should be a preclinical assessment of cardiac ion channels inhibition and QT interval (i.e. time duration between the start of the Q wave and the end of the T wave on an electrocardiogram) prolongation caused by small compounds[8]. This is defined as cardiotoxicity and involves the inhibition of any of the three cardiac ion channels: the voltage-gated potassium channel (hERG), the voltage-gated calcium channel (Cav1.2), and the voltage-gated sodium channel (Nav1.5). The pharmaceutical industry suffers significant losses due to cardiotoxicities that emerge during the early, preclinical, or clinical stages of drug development, leading to the withdrawal of several drugs from the market and the halting of many drug discovery programs in their pipelines [9]. Examples of such drugs are astemizole, terfenadine, sertindole, grepafloxacin, vardenafil, cisapride, and ziprasidone, which have been withdrawn or severely restricted on the use for the undesirable cardiac toxicity side effects [10][11][12].

An effective solution that offers a prominent alternative to reduce costs and advance the development of lead candidates is the field of computer-aided drug discovery (CADD) [6][13][14]. Toxicity prediction algorithms have recently become a very important component of modern CADD [13], with cardiotoxicity prediction algorithms as the most dominant of these methods.

In recent years, researchers have increasingly utilized machine learning (ML) algorithms to construct and deploy robust models for predicting cardiac ion channels’ inhibition. Previous reviews [10] [15] have indicated that during the early 2000s, statistical approaches such as Naïve Bayes, Gaussian processes, expectation–maximization, and partial least squares (PLS) were commonly employed ML algorithms [16][17][18][19][20]. The majority of those published prediction methods used less than a thousand compounds to train their models. Later, there has been a notable shift toward the utilization of random forest (RF), support vector machine (SVM), and deep neural network (DNN) methods [21][22][23][24], using up to 15,000 compounds for training the models. This transition is primarily due to their superior empirical performance in the field, as evidenced by multiple research publications outlined recently [25].

However, there is still important room for improvement of cardiotoxicity prediction. First, almost all published methods during the last 20 years focused only on hERG liability prediction, as there were very few bio-activity data available on the other two cardiac ion channels. Second, different models were trained on different datasets lacking a common and unique dataset to use for benchmarking. Even though most papers used similar data sources, different curation techniques and decisions were applied. Third, published and evaluated models by some authors cannot be validated for better generalization as many published models show overlap between training and test sets; hence presenting over-optimistic performance results. Illustrative examples include the work of Konda et al. [26], wherein their test set exhibits an approximate 50% overlap in molecular compounds with the training set. This dataset has been extensively employed and cited by over 20 other scholarly works. Another noteworthy scenario is presented by Liu et al. [27], who introduced a model trained and assessed on the dataset published by Doddareddy et al. [28], which itself has garnered over 100 citations. Liu et al.’s findings indicated that although their best model demonstrated enhanced performance on the provided test set (achieving an accuracy of 88% for [28] versus 91% for [27]), its performance declined when evaluated on external sets, reaching 47% and 58% respectively. Our own evaluation further revealed that more than a quarter of the test data exhibited an overlap with the training set, exceeding 80% in terms of structural similarity.

In this study, we present a framework to perform further analyses on a very large open-access unique and comprehensive hERG, Nav1.5, and Cav1.2 cardiotoxicity integrated database of small molecules and their activities. We also present a deep learning model used to benchmark different feature sets, namely descriptors, fingerprints, and graph-based representations, for cardiotoxicity prediction. Finally, we benchmark our best models on strictly unique extracted external evaluation sets and provide a robust predictive model for each target.

## 2. Methods

### 2.1. Compilation of a Cardiotoxicity Database

The compounds demonstrating inhibitory activity used in this study were sourced from various public data repositories, including the ChEMBL bioactivity database [29][30][31], PubChem [32], BindingDB [33][34], hERGCentral [35], and US patent and literature-derived data [36][37][38][39]. The collected data was split into two primary classes based on the available information: IC50-type and inhibition-type values. The IC50-type measurements encompassed inhibitory activity values expressed as half maximum inhibitory concentration (IC50), half maximum effective concentration (EC50), median effective dose (ED50), inhibitory constant (Ki), or dissociation constant (Kd). Conversely, the inhibition-type measurements comprised percentage inhibition values at specific concentrations. In the case of inhibition-type entries, a manual examination of assay descriptions was conducted to extract the relevant activity values. Values reported with thresholds other than 50% inhibition (e.g., IC70, IC30, IC20, etc.), raw measurements of current, ratios indicating prolonged QT intervals, and similar values were excluded. This procedure aligns with the methodology outlined by Sato et al. [40]. Additionally, for hERG data, cross-referencing with the hERGCentral retrieved data was performed, and any erroneous activity values were rectified.

Following this step, the chemical structures within each dataset underwent standardization using the Python packages RDKit (http://www.rdkit.org) and MolVS (https://github.com/mcs07/MolVS). The standardization procedure encompassed selecting the largest fragment, eliminating explicit hydrogens, ionization, and calculating of stereochemistry. Next, the compounds were encoded as SMILES (Simplified Molecular Input Line Entry System) strings. Because multiple SMILES strings can correspond to the same structure, and there is no universal approach for generating a canonical SMILES string, we converted the SMILES strings to InChI (International Chemical Identifier) keys [41] using RDKit to identify duplicate compounds. All potency values were transformed into nanomolar (nM) units. For duplicate compounds, the mean value was subsequently computed, while only retaining values falling within the 95% quantile range to exclude excessive outliers. The overall methodology employed to construct the extensive small molecule cardiotoxicity database is illustrated in Figure 1.

**Figure 1.**
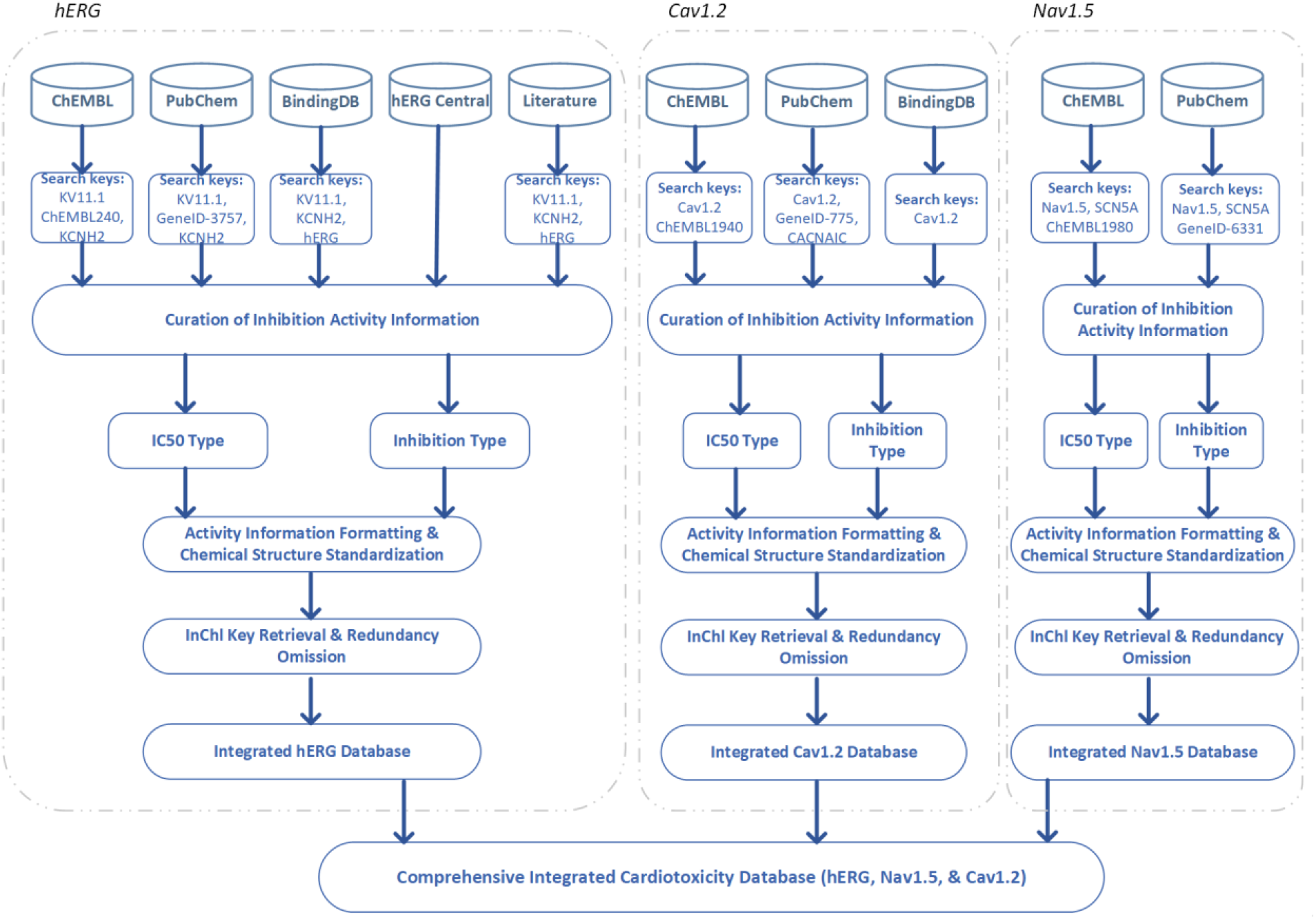
Schematic of the gathering, curation, and integration pipeline to generate the comprehensive integrated cardiotoxicity database of small molecules for all three ion channels: hERG, Nav1.5, and Cav1.2.

All activities were then transformed into molar values and subsequently standardized by calculating the pIC50 as follows :

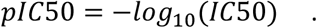

Compounds were classified based on their IC50 values, following standard criteria used by researchers in the field [25][40][42]. Compounds with IC50 values of 10 μM or below (pIC50 ≥ 5) were categorized as blockers (inhibitors), while compounds with IC50 values higher than 10 μM (pIC50 < 5) were categorized as non-blockers (inactive).

In adherence to data science best practices for model development, as illustrated in Figure 2, we extracted two external/independent test sets from each target dataset (hERG, Nav1.5, and Cav1.2). We used RDKit (default settings) to compute the Tanimoto similarity [43][44] between each pair of compounds in the datasets using 2048-bit extended connectivity fingerprints, also referred to as circular or Morgan fingerprints [45]. The first test set consisted of compounds with a structural similarity of no more than 60% (Tanimoto similarity ≤ 0.6) to the remaining development set, while the second test set comprised compounds with a structural similarity of no more than 70% (Tanimoto similarity ≤ 0.7) to the remaining development set. These external unique sets were denoted as hERG-70 & hERG-60 for hERG, Nav-70 & Nav-60 for Nav1.5, and Cav-70 & Cav-60 for Cav1.2. The composition of each set as derived from the upstream data sources can be observed in supplementary figure S1. Our unique development sets were subsequently partitioned into training and validation sets using an 80/20 ratio for each target. These splits were used for hyperparameter tuning and were stratified by pIC50. Note that random splitting of the training and validation sets can lead to splits that share portions of similar data points, which in turn, can lead to a positive bias in validation performance compared to our external test sets. However, any bias in validation performance is minimal due to multiple factors: the development set of curated molecular compounds that is already screened for distinct and unique structures, the stratification strategy, the size of the validation set, and the biological nature of our targets, namely, hERG, Nav1.5, and Cav1.2. These cardiac ion channels possess promiscuous binding sites at their central cavity that can interact with a diverse set of chemical structures [46][47][48]. In other words, hERG, Nav1.5, and Cav1.2 are not similar to an enzyme or a receptor with a specific catalytic site. Enzyme targets can usually interact with a distinct set of compounds with a common chemical scaffold. Given this additional biological parameter, which adds an intrinsic diversifying factor to our dataset, we believe that the derived validation and training datasets will already be diverse by nature.

**Figure 2.**
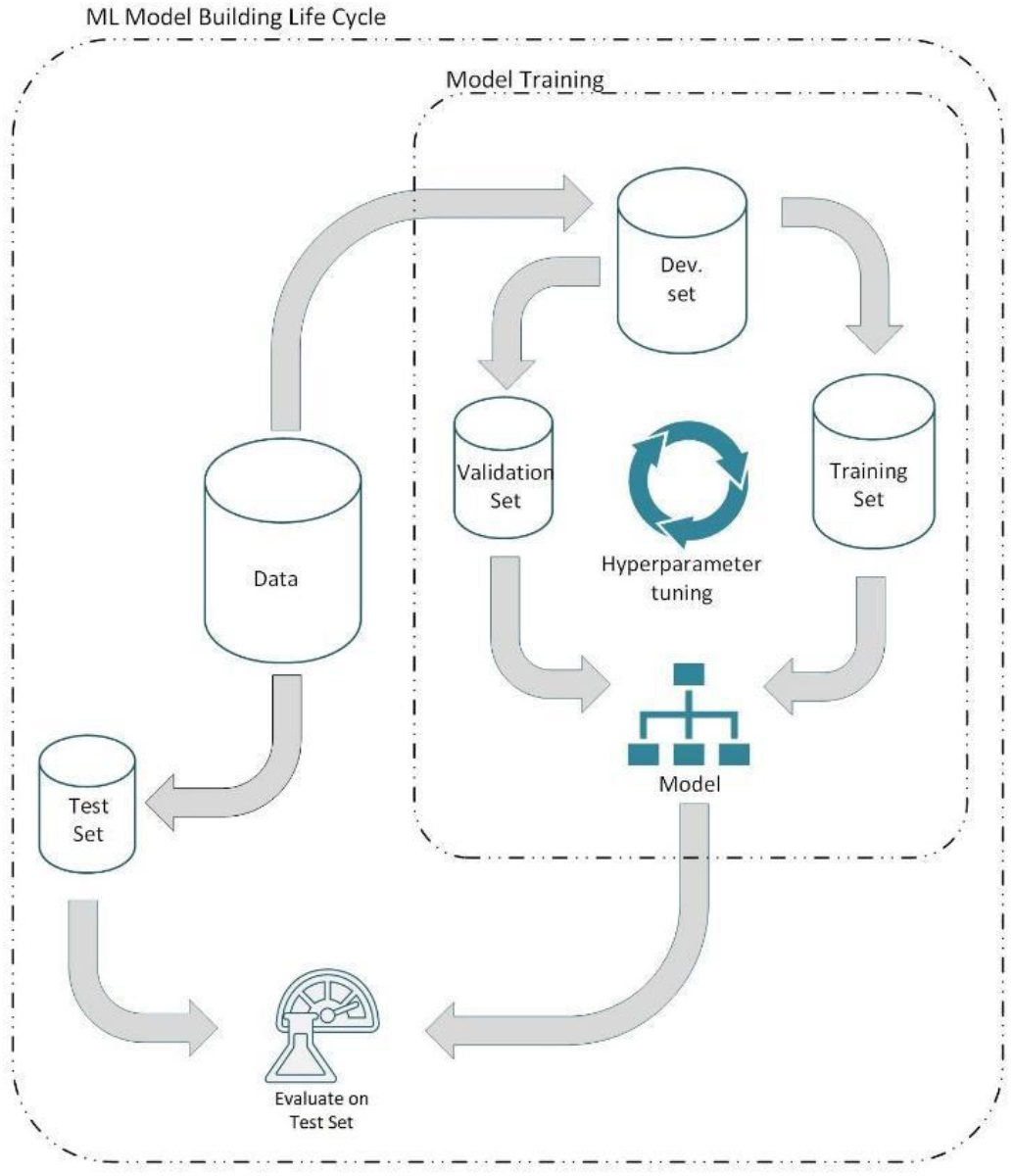
Data science methodology for hyperparameter tuning, monitoring overfitting, and best model selection.

### 2.2. Molecular Features

The chemical structures in our database are represented in the SMILES format and serve as the primary input for our predictive model training. Prior to conducting analyses, these structures need to be transformed into numerical representations. In this research, we explored three types of feature representations: molecular fingerprints, molecular descriptors, and molecular graph representations.

The PyBioMed Python package [49] was used to compute molecular fingerprints. This process generated two types of fingerprints: extended connectivity fingerprints with a maximum diameter parameter of two (ECFP2) (a vector of 1024 ECFP fingerprint values) and PubChem fingerprints (a vector of 881 values).

The calculation of molecular descriptors was performed using the Mordred Python package [50]. We employed 2D descriptors only as these require fewer computational resources compared to 3D descriptors without sacrificing predictive performance [14][51][52], resulting in a total of 1613 descriptors. Preprocessing and feature selection were accomplished through a Scikit-Learn [53] pipeline consisting of four modules. First, a univariate imputer was employed to discard columns with no calculated values and replace missing values in other columns with the mean. Second, a standardization step was applied to remove the mean and scale the values to unit variance. Third, zero-variance features were removed. Finally, for any pair of highly correlated features (Pearson correlation above 0.95), one of them was randomly discarded, as such correlated features convey nearly identical information. Consequently, the preprocessing procedure reduced the feature set for each target database (i.e. hERG, Nav1.15, and Cav1.2) to 806, 549, and 681 descriptors, respectively (refer to Tables S1, S2, and S3 of the supplementary data for the respective descriptor names).

Although SMILES are a convenient linear textual representation of molecules, a graph representation is closer to reality. Molecules can be represented as graphs, wherein atoms are depicted as nodes and chemical bonds between atoms are depicted as edges. These two elements, nodes and edges, constitute the fundamental components of a graph and can serve as a molecular graph representation to encode the topological information inherent within molecules. To clarify more, this featurization process of a molecular compound yields two data structures. First, each atom/node is represented by a vector that can encode any type of information describing the atom. This is called the nodes embedding matrix, which has a dimension of M × k, with M the number of atoms in the molecule and k the number of node features. Here, we used 67 different node features according to the previously described node featurization strategy by Ryu et al. [55], including features such as the atom symbol, node degree, number of bound hydrogens, implicit valence, aromaticity, and size of the ring containing the atom, and further detailed in the supplementary table S4. As illustrated in Figures S2, S3, and S4, the value of M ranges up to 122 for hERG, 294 for Nav1.5, and 87 for Cav1.2. Second, the bonds/edges are encoded in an adjacency matrix that indicates the bonds between atoms [54]. As constrained by the deep learning model we used (see below for details), the adjacency matrix of size M × M is converted to an edge list of dimensions 2N × 2, with N the number of edges. RDKit was used to compute the node features and the edge list of the molecular structures.

### 2.3. Model Architecture

A neural network architecture was designed to accommodate the different features described above within a unified machine learning model. This system offers flexibility, allowing for the addition, removal, enabling, or disabling of different modules within the network. This flexibility is particularly advantageous during hyperparameter tuning. Figure 3 presents a simplified schematic of the architecture employed in this research.

**Figure 3.**
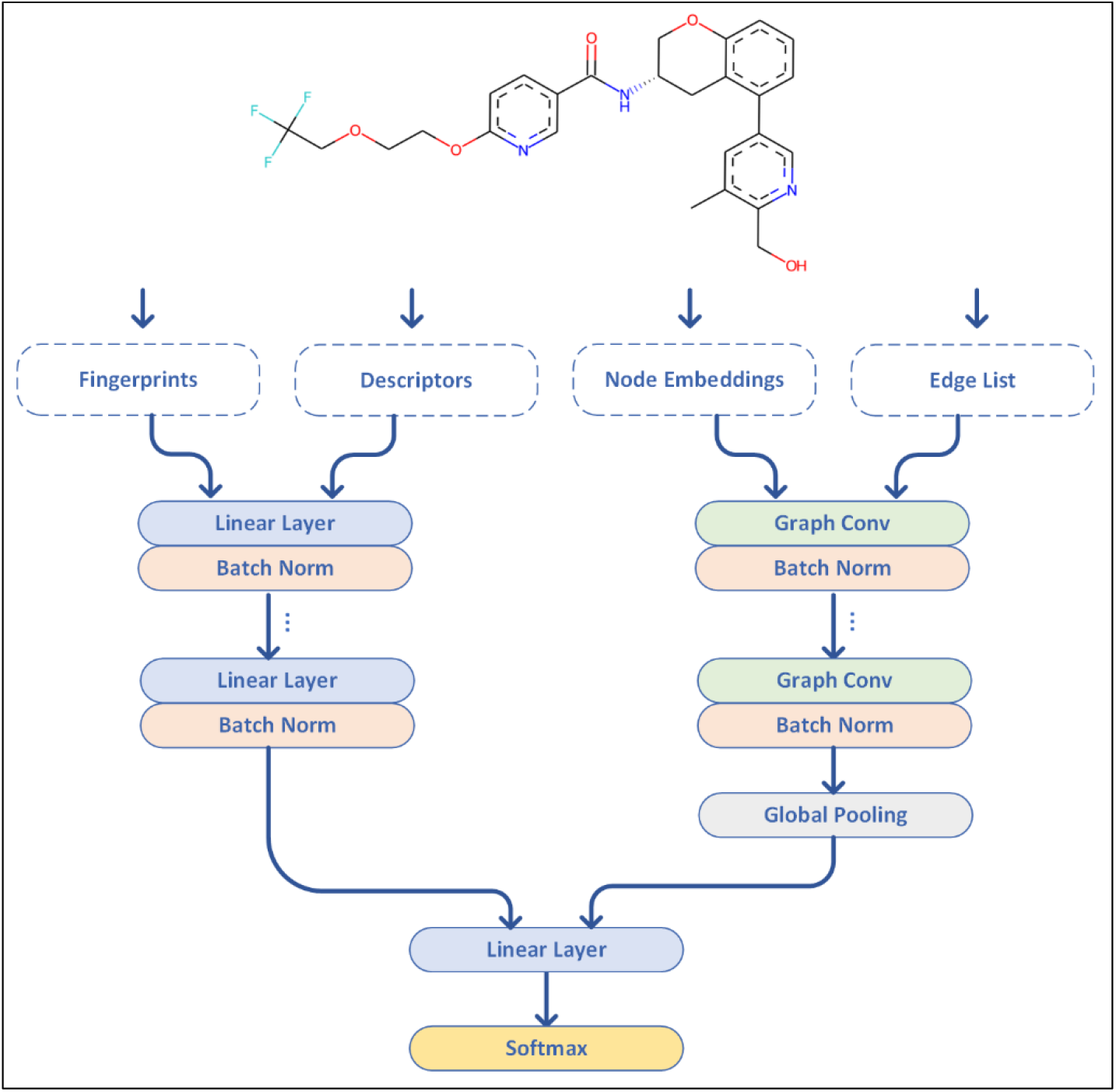
Simplified conceptual visualization of the deep learning model used in this study. The architecture can enable/disable any path, layer, or feature set of the molecule.

The architecture consists of two paths that are merged at the final layer. The first path accepts descriptors, fingerprints, or a combination of both feature vectors as input. The vector is then passed through a series of sequential blocks, with each block comprising a linear layer followed by batch normalization [56]. The second path takes the graph representation of the chemical structure as input. This graph representation is composed of a node embedding matrix with dimensions M × k and an edge list represented as a 2D matrix with dimensions 2N×2. These matrices undergo a sequence of feature extraction blocks, with each block consisting of a graph convolutional layer (GCN) [57] followed by batch normalization. For batch processing of chemical structures, zero padding was employed. Following the graph convolutions, a global pooling layer is applied to reduce the dimensionality of the generated tensor to a 1D vector. This vector is then concatenated with the intermediate representations that capture higher-level abstractions within the input chemical structure, as encoded by either the fingerprints or descriptors. The final output layer utilizes a softmax layer, predicting whether the compound is an inhibitor or not. Hyperparameter tuning was employed to find the best set of hyperparameters from a predefined dictionary as summarized in Supplementary Table S5.

### 2.4. Evaluation Metrics

Initially, all models underwent evaluation on validation sets stratified by pIC50 derived from the development sets, consisting of 20% of the data. The best-performing hyperparameters for each combination of feature representations were identified based on their performance on the validation set. These hyperparameters were then used to construct the final best models, which were subsequently evaluated on external test sets comprising 60% and 70% structurally dissimilar molecular compounds.

The performance of the models was assessed using multiple binary evaluation metrics, including accuracy (AC), sensitivity (SE), specificity (SP), F1-score (F1), correct classification rate (CCR), and Matthew’s correlation coefficient (MCC). Accuracy represents the overall predictive effectiveness of a classifier, while sensitivity and specificity measure the predictive powers for positive and negative instances, respectively. The CCR quantifies the proportion of instances that are correctly classified by the model. The F1-score computes the harmonic mean of precision and sensitivity. The MCC takes into account the balance ratios of the four categories in the confusion matrix (TP, TN, FP, FN) and provides an objective reflection of the model’s predictive power. The final model selection was performed based on the F1 score.

The definitions of these evaluation metrics are provided as follows:

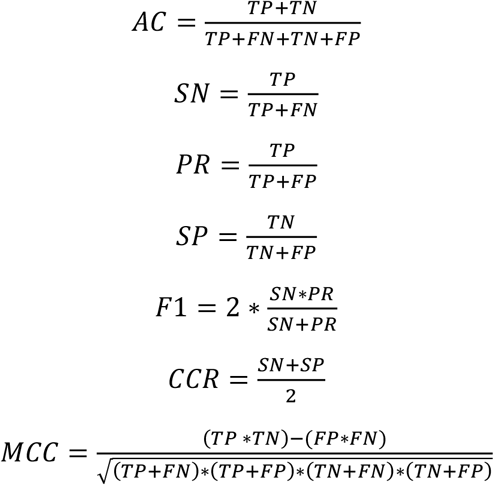

### 2.5. hERG benchmarked tools settings

#### 2.5.1. CardioTox

CardioTox [21] is a command line-based prediction tool for hERG inhibition. The method uses an ensemble strategy of five models each trained on: 2D+3D descriptors, molecular graph representation, molecular fingerprints, SMILES embedding, and a fingerprint embedding model. The outputs of each model are concatenated, after which the final binary prediction (hERG blocker or non-blocker) is made. For benchmarking purposes we used the default settings as outlined in the corresponding GitHub repository (https://github.com/Abdulk084/CardioTox).

#### 2.5.2. CardPred

CardPred [58] is a recent web-based prediction tool for hERG inhibition (http://ssbio.cau.ac.kr/CardPred). The method uses a deep learning network trained on a set of 2130 molecular compounds. The predictions are based on a combination of descriptors and fingerprints calculated using an external software, DRAGON (version 7.0.10) [59]. The tool predicts the probability of the chemical structure being a blocker or not. We used the default settings and a threshold of greater or equal than 50% to decide on hERG blockers in this study.

#### 2.5.3. ADMETsar 2.0

ADMETsar 2.0 [60] is an optimized version of the previously released version of ADMETsar. It is a comprehensive open source and free tool for the prediction of different chemical ADMET properties. The web-based (http://lmmd.ecust.edu.cn/admetsar2/) prediction tool takes as input SMILES strings and comprises 47 different models for drug discovery, among them the hERG cardiotoxicity prediction model. Default settings were used to benchmark this tool.

#### 2.5.4. ADMETlab 2.0

ADMETlab 2.0 [61] is another comprehensive web server (https://admetmesh.scbdd.com/) for the predictions of pharmacokinetics and toxicity properties of chemicals using a multi-task graph attention framework. The web-based prediction tool takes as input SMILES strings to make multiple predictions, including hERG cardiotoxicity predictions. Default settings were used to benchmark this tool.

### 2.6. Code availability

CToxPred, a comprehensive cardiotoxicity prediction method, is available as an open-source Python command-line tool and can be called from a notebook. It uses Pybel (version 0.13.2) [62], Open Babel (version 3.1.1) [63], and PyBioMed (version 1.0) [49] to compute PubChem & ECFP2 fingerprints; Mordred (version 1.2.0) [50] to calculate molecular descriptors; RDKit (version 2022.09.1) [64] for chemical structure information retrieval, used for example in the graph representation construction and other tasks; Scikit-Learn (version 1.0.2) [53] for pipeline data preprocessing and evaluation metric calculations; PyTorch (version 1.12.1) & PyTorch Geometric (version 2.3.1) [65] libraries to implement our deep learning and graph neural network architecture; and NumPy (version 1.21.6) [66], SciPy (version 1.7.3) [67], and Pandas (version 1.3.5) [68] for scientific computing. Matplotlib (version 3.5.3) [69] and Seaborn (version 0.12.2) [70] were used for data visualization. Data analysis was performed using Jupyter notebooks [71].

CToxPred is available as open source under the permissive MIT license on GitHub at https://github.com/issararab/CToxPred. Analysis notebooks to reproduce the presented results are also available in the same repository.

## 3. Results

### 3.1. A Comprehensive Database of Cardiac Ion Channel Blockers

The presented collection of data establishes a framework intended for researchers operating within the realm of drug discovery to conduct in-depth analyses and further studies. This collection includes a large and freely accessible unique and comprehensive hERG, Nav1.5, and Cav1.2 cardiotoxicity integrated database of small molecules and their activities. The database was sourced from a variety of public repositories, such as databases like ChEMBL, PubChem, BindingDB, and hERGCentral, as well as from US patent data and literature mining. Figure 4 shows a quick overview of the data composition for each target (hERG, Nav1.5, and Cav1.2), indicating the sources from which it was gathered.

**Figure 4:**
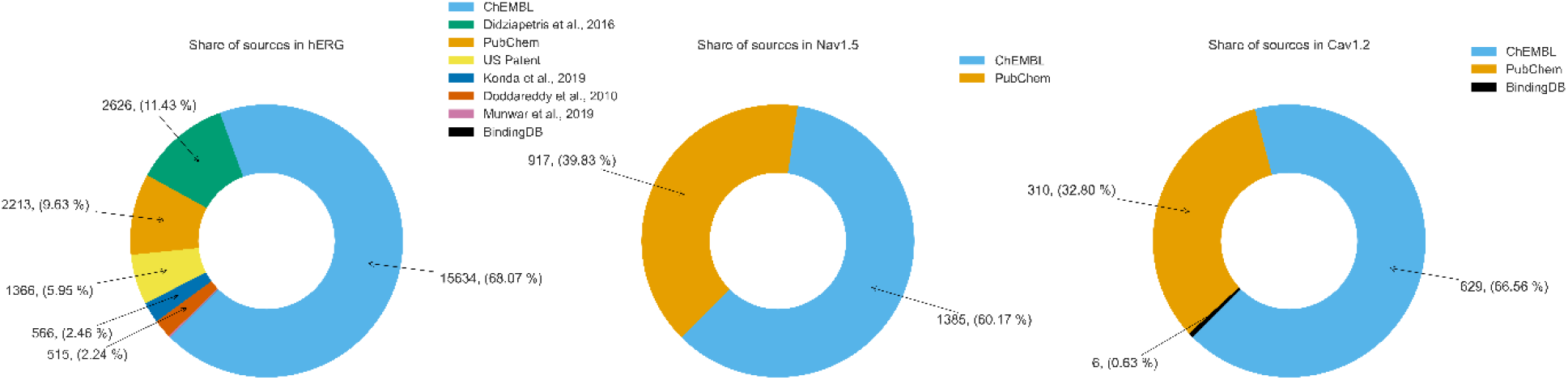
The composition of each target (hERG, Nav1.5, and Cav1.2) as derived from the upstream data sources in the final extensive cardiotoxicity database.

External test sets were derived from the collected database using two Tanimoto similarity thresholds, namely 60% and 70% structural similarity, as illustrated in the pairwise Tanimoto similarity distributions in Figure 5. These sets were extracted in a manner that ensures the preservation of the pIC50 distribution found in the development set. This approach ensures that the blocker vs. non-blocker class distribution is maintained within each set, thereby enabling reliable performance evaluation of the built models. Figure 6 illustrates the pIC50 density distribution in each dataset, which also supports the community choice of 10μM as a threshold for discriminating between blockers and non-blockers. The distribution of compounds in each class exhibits a ratio of approximately 6:4, favoring blockers, for each respective set, as illustrated in Table 1.

**Table 1.** Class distribution of blockers vs non-blockers in each set and for each target ion channel.

|  | hERG |  | Nav1.5 |  | Cav1.2 |  |
| --- | --- | --- | --- | --- | --- | --- |
| Property | Blockers | Non-Blockers | Blockers | Non-Blockers | Blockers | Non-Blockers |
| Dev. Set | 13428 | 8818 | 1423 | 673 | 530 | 272 |
| Eval-70 | 264 | 209 | 97 | 45 | 52 | 29 |
| Eval-60 | 125 | 125 | 48 | 16 | 41 | 21 |

**Figure 5.**
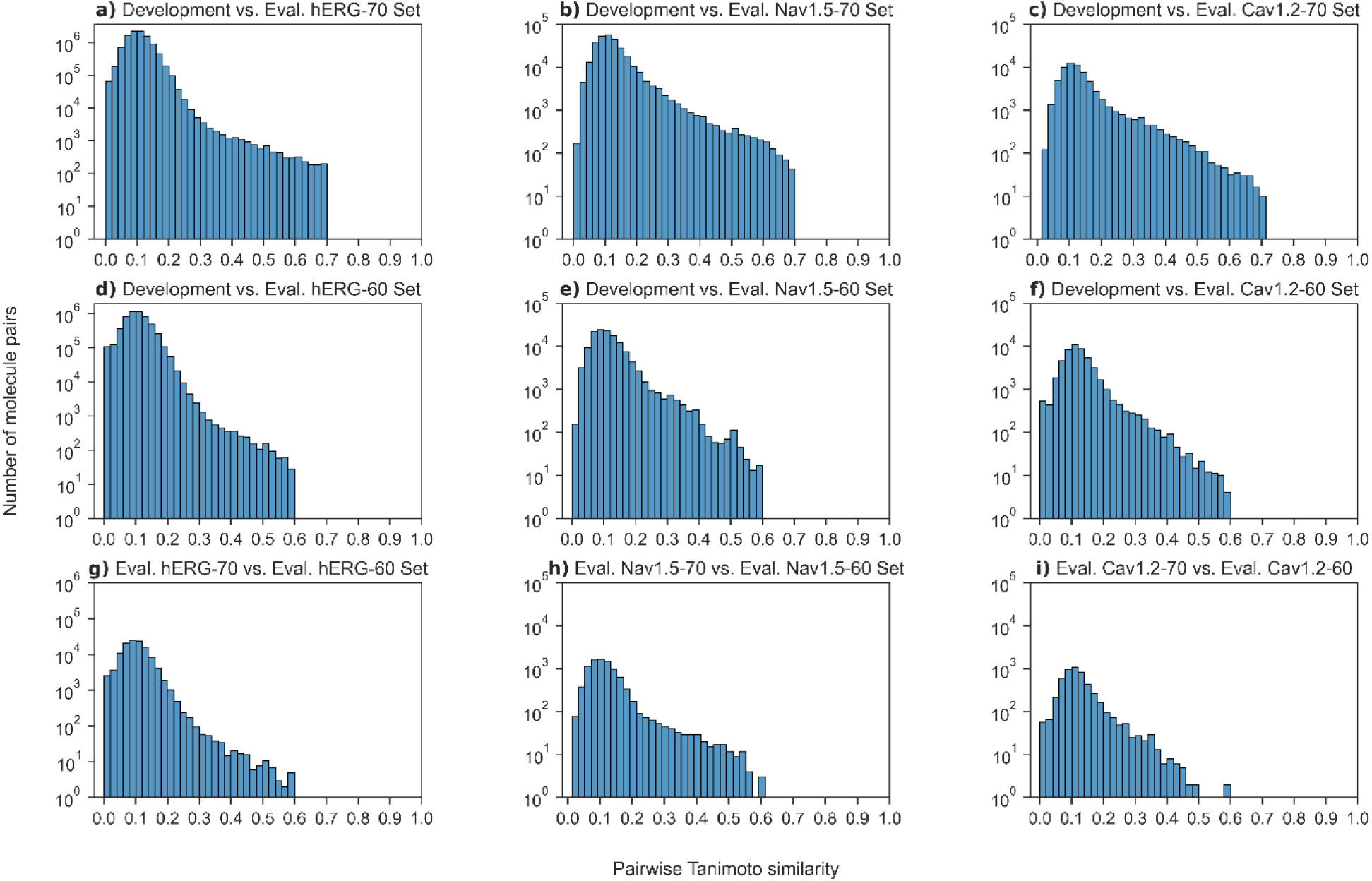
Distribution of the pairwise Tanimoto similarity for each molecule in the **(a)** hERG development set with the ones in the evaluation set hERG-70, **(b)** Nav1.5 development set with the ones in the evaluation set Nav-70, **(c)** Cav1.2 development set with the ones in the evaluation set Cav-70, **(d)** hERG development set with the ones in the evaluation set hERG-60, **(e)** Nav1.5 development set with the ones in the evaluation set Nav-60, **(f)** Cav1.2 development set with the ones in the evaluation set Cav-60, **(g)** hERG-70 evaluation set with the ones in the evaluation set hERG-60, **(h)** Nav-70 evaluation set with the ones in the evaluation set Nav-60, and **(i)** the Cav-70 evaluation set with the ones in the evaluation set Cav-60.

**Figure 6.**
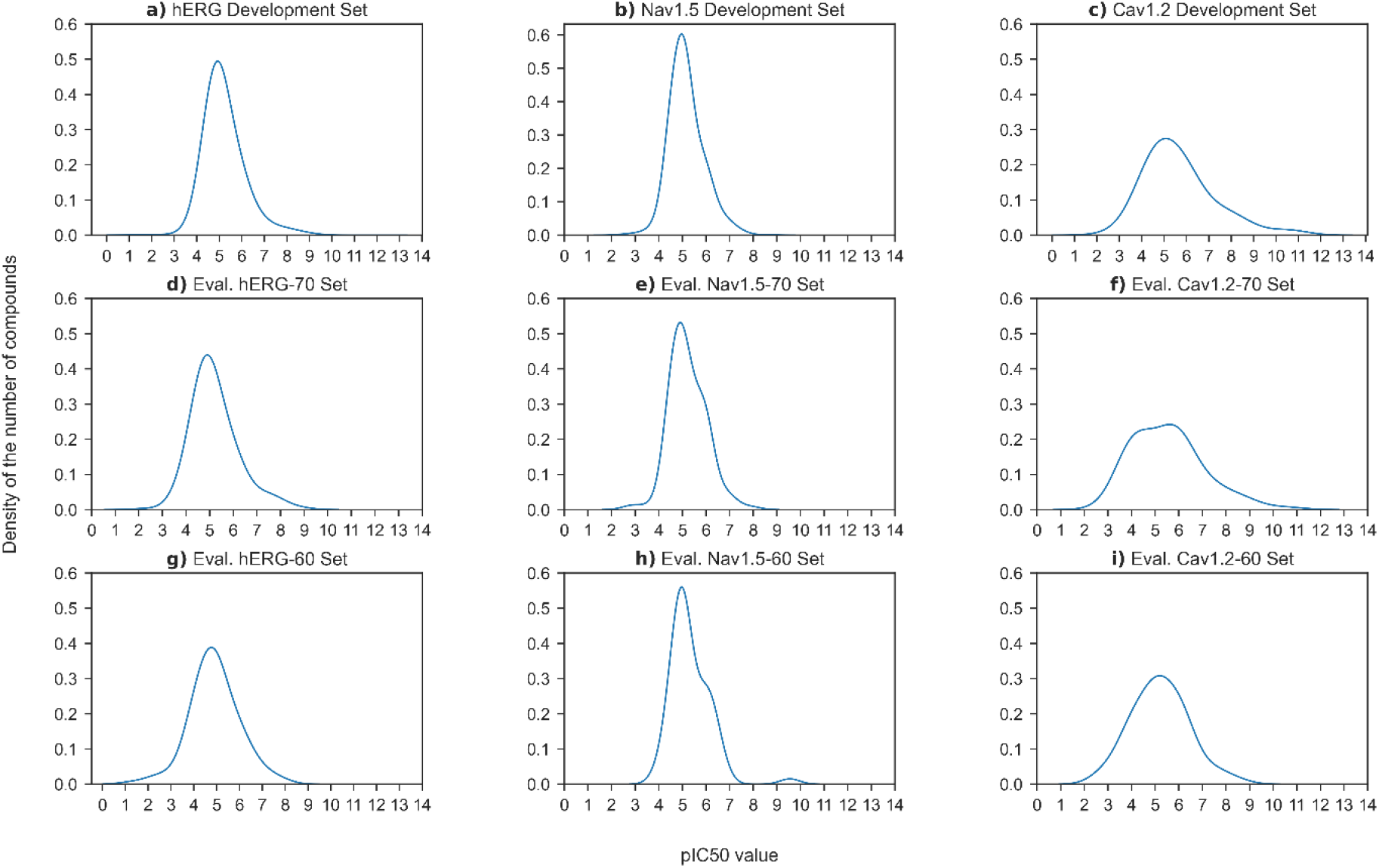
Density distribution of the pIC50 activity of molecular compounds in the **(a)** hERG development set, **(b)** Nav1.5 development set, **(c)** Cav1.2 development set, **(d)** hERG-70 evaluation set, **(e)** Nav-70 evaluation set, **(f)** Cav-70 evaluation set, **(g)** hERG-60 evaluation set, **(h)** Nav-60 evaluation set, and **(i)** Cav-60 evaluation set.

### 3.2. Evaluation of Molecular Features For Machine Learning of Cardiotoxicity

The deep learning model presented in this study has been designed to take different combinations of feature representations of a molecular structure as input. For each target’s (i.e. hERG, Nav1.5, Cav1.2) molecular compounds dataset, the network was trained on seven combinations from the three feature sets (fingerprint, descriptor, and graph representation). Conducting a thorough exploration of the hyperparameter search space, as detailed in Table S5, yields a total of 576 distinct combinations for each feature combination. In order to mitigate the computational cost associated with this exhaustive procedure, a random search policy [72] was employed, involving only 100 randomly selected hyperparameter combinations. The final optimal set of hyperparameters was determined based on optimal performance on the validation set, which accounts for 20% of the development set. The training process was performed on an NVIDIA T4 with 16GB dedicated RAM, granted by Google Cloud Research. For each feature combination, the best-performing model was selected and subsequently evaluated on our two external datasets. We name the best models CToxPred-hERG, CToxPred-Nav, and CToxPred-Cav for each of the targets respectively.

#### 3.2.1. hERG Cardiotoxicity Prediction

We assessed the optimal model for each combination of molecular features on two separate external test sets to evaluate hERG cardiotoxicity predictions. Surprisingly, the results revealed that standard fingerprints outperformed more complex feature representations. Nonetheless, it is worth noting that the combination of fingerprints with additional complex features showcased competitive performance, potentially enhancing generalization. This trend was demonstrated by the competitive performance of the combined features on hERG-60 versus hERG-70 datasets. However, the more different the test data is from the training data, the lower the performance, as evidenced by the drop in performance from the hERG-70 to hERG-60 test sets (an observation that was expected). For additional observations and insights, please consult Table 2 and Figure S8.

**Table 2.**
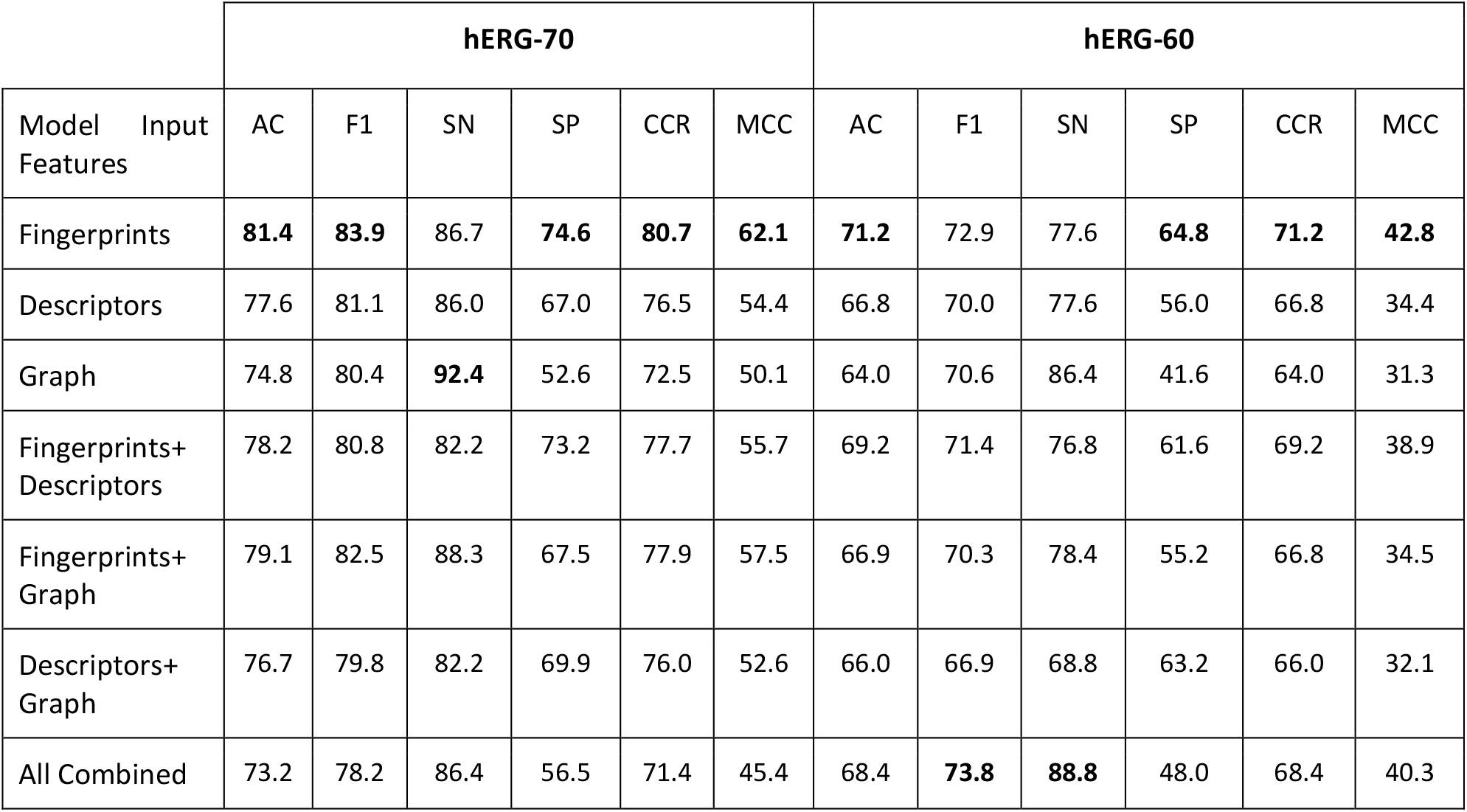
hERG toxicity prediction performance of each feature combination using the deep learning model on the two external test sets hERG-70 and hERG-60.

| Model Input Features | hERG-70 |  |  |  |  |  | hERG-60 |  |  |  |  |  |
| --- | --- | --- | --- | --- | --- | --- | --- | --- | --- | --- | --- | --- |
|  | AC | F1 | SN | SP | CCR | MCC | AC | F1 | SN | SP | CCR | MCC |
| Fingerprints | **81.4** | **83.9** | 86.7 | **74.6** | **80.7** | **62.1** | **71.2** | 72.9 | 77.6 | **64.8** | **71.2** | **42.8** |
| Descriptors | 77.6 | 81.1 | 86.0 | 67.0 | 76.5 | 54.4 | 66.8 | 70.0 | 77.6 | 56.0 | 66.8 | 34.4 |
| Graph | 74.8 | 80.4 | **92.4** | 52.6 | 72.5 | 50.1 | 64.0 | 70.6 | 86.4 | 41.6 | 64.0 | 31.3 |
| Fingerprints+ Descriptors | 78.2 | 80.8 | 82.2 | 73.2 | 77.7 | 55.7 | 69.2 | 71.4 | 76.8 | 61.6 | 69.2 | 38.9 |
| Fingerprints+ Graph | 79.1 | 82.5 | 88.3 | 67.5 | 77.9 | 57.5 | 66.9 | 70.3 | 78.4 | 55.2 | 66.8 | 34.5 |
| Descriptors+ Graph | 76.7 | 79.8 | 82.2 | 69.9 | 76.0 | 52.6 | 66.0 | 66.9 | 68.8 | 63.2 | 66.0 | 32.1 |
| All Combined | 73.2 | 78.2 | 86.4 | 56.5 | 71.4 | 45.4 | 68.4 | **73.8** | **88.8** | 48.0 | 68.4 | 40.3 |

Considering all evaluation metrics, we can rank the individual feature sets in terms of performance from best to worst as follows: fingerprints, descriptors, and then graph representations. The fingerprint-based model exhibited the highest accuracy and F1-score for both evaluation sets. The F1-score for hERG-60 demonstrated a comparative value to the models that utilized all combined features. Therefore, we have selected the fingerprint-based model as our best-performing model for predicting hERG cardiotoxicity of small molecules. Throughout the rest of the analysis, we will refer to this model as CToxPred-hERG, which has an average standard error of ±1.8%. For the confusion matrices of CToxPred-hERG on the two external test sets, please refer to Supplementary Figure S8.

We also conducted a comprehensive benchmarking study of CToxPred-hERG, comparing it with another published command-line tool called CardioTox [21], as well as three web-based prediction tools: CardPred [58], ADMETsar 2.0 [60], and ADMETlab 2.0 [61]. All models were evaluated using the same external sets of molecular compounds (Table 3). As anticipated, all models demonstrated superior performance on the hERG-70 dataset and relatively lower performance on the hERG-60 dataset.

**Table 3.** Performance evaluation of CToxPred-hERG compared to CardPred, ADMETsar, ADMETlab, and CardioTox on the 2 external test sets.

|  | hERG-70 |  |  |  |  |  | hERG-60 |  |  |  |  |  |
| --- | --- | --- | --- | --- | --- | --- | --- | --- | --- | --- | --- | --- |
| Model | AC | F1 | SN | SP | CCR | MCC | AC | F1 | SN | SP | CCR | MCC |
| CardPred | 56.1 | 57.0 | 52.7 | 60.3 | 56.5 | 13.0 | 53.9 | 45.4 | 37.9 | 70.2 | 54.1 | 8.6 |
| ADMETsar 2.0 | 68.5 | 75.0 | 84.5 | 48.3 | 66.4 | 35.5 | 66.4 | 70.6 | 80.8 | 52.0 | 66.4 | 34.3 |
| ADMETlab 2.0 | 71.7 | 73.8 | 71.6 | 71.8 | 71.7 | 43.1 | 68.0 | 67.5 | 66.4 | 69.6 | 68.0 | 36.0 |
| CardioTox | 81.2 | 83.1 | 83.0 | **78.9** | **81.0** | 61.9 | **80.4** | **78.4** | 71.2 | **89.6** | **80.4** | **61.9** |
| CToxPred-hERG | **81.4** | **83.9** | **86.7** | 74.6 | 80.7 | **62.1** | 71.2 | 72.9 | **77.6** | 64.8 | 71.2 | 42.8 |

Among the competing models, CardPred exhibited the lowest performance, while CardioTox and CToxPred-hERG displayed comparable results on the hERG-70 dataset. However, CardioTox outperformed our best model on the hERG-60 dataset. Two hypotheses can be inferred from these findings: either CardioTox can generalize better to unknown molecules, or the evaluation data used for CardioTox included compounds that were already present in its training data. Further investigations revealed that the training data used to construct the CardioTox model had a 40% overlap with the hERG-70 dataset and a 68% overlap with the hERG-60 dataset. As such, we anticipate that there is a positive bias in these evaluation results for CardioTox. In order to conduct an unbiased comparison between the two models, it would be necessary for both methods to be trained on the same data and evaluated using the same test set.

#### 3.2.2. Nav1.5 Cardiotoxicity Prediction

Similar to the evaluation performed for hERG, we also assessed the most effective predictive model for Nav1.5 cardiotoxicity using various combinations of molecular features on the two external test sets. For the Nav-70 test set, three feature combinations demonstrated superior and comparable performance in terms of both F1-score and AC. These combinations were *‘fingerprints + descriptors’, ‘fingerprints + graph representation’*, and *‘all combined’* features. Thus, we can infer that, similar to hERG, fingerprints provide more informative representations to discriminate blocksers from non-blockers. This conclusion is further supported by the superior performance exhibited by all features in the hERG-70 set. However, it is worth noting that this observation does not hold for structurally dissimilar compounds, as demonstrated in the Nav-60 evaluation, where descriptors performed better (as shown in Table 4 and Figure S10).

**Table 4.** Nav1.5 toxicity prediction performance of each feature combination using the deep learning model on the two external test sets Nav-70 and Nav-60.

| Model Input Features | Nav-70 |  |  |  |  |  | Nav-60 |  |  |  |  |  |
| --- | --- | --- | --- | --- | --- | --- | --- | --- | --- | --- | --- | --- |
|  | AC | F1 | SN | SP | CCR | MCC | AC | F1 | SN | SP | CCR | MCC |
| Fingerprints | 80.3 | 85.3 | 83.5 | 73.3 | 78.4 | 55.6 | 62.5 | 72.7 | 66.7 | 50.0 | 58.3 | 14.9 |
| Descriptors | 79.6 | 84.7 | 82.5 | 73.3 | 77.9 | 54.4 | 75.0 | 83.0 | 81.2 | 56.2 | 68.8 | 36.1 |
| Graph | 77.5 | 83.7 | 84.5 | 62.2 | 73.4 | 47.3 | 70.3 | 78.7 | 72.9 | **62.5** | 67.7 | 32.0 |
| Fingerprints + Descriptors | **81.7** | **86.5** | **85.6** | 73.3 | 79.5 | 58.2 | **76.6** | **84.2** | **83.3** | 56.2 | **69.8** | **38.8** |
| Fingerprints + Graph | **81.7** | **86.5** | **85.6** | 73.3 | 79.5 | 58.2 | 60.9 | 72.5 | 68.8 | 37.5 | 53.1 | 5.8 |
| Descriptors + Graph | 78.2 | 83.4 | 80.4 | 73.3 | 76.9 | 51.9 | 71.9 | 80.9 | 79.2 | 50.0 | 64.6 | 28.1 |
| All Combined | **81.7** | 86.2 | 83.5 | **77.8** | **80.6** | **59.4** | 71.9 | 81.6 | **83.3** | 37.5 | 60.4 | 21.8 |

Considering different evaluation metrics and individual feature sets, we can rank the best-performing features from worst to best as follows: fingerprints, graphs, and then molecular descriptor representations. Although the ranking order may vary when considering each test set individually, we tend to prioritize the Nav-60 set as it is structurally dissimilar and therefore represents better generalization to unseen data. Interestingly, the combination of fingerprints and descriptors exhibited the highest performance across most metrics and on both test sets. This combination can be seen as an enrichment of complementary feature information, where one set excels at predicting Nav-70 compounds while the other performs better for Nav-60 molecular structures.

Based on these findings, the model that combines fingerprints and descriptors was selected as the optimal model for the Nav1.5 cardiotoxicity prediction task and named CToxPred-Nav1.5. It exhibited an average standard error of ±3.2%. For the confusion matrices detailing the predictions of CToxPred-Nav1.5 on the two external test sets, please refer to the SOM, Figure S11.

#### 3.2.3. Cav1.2 Cardiotoxicity Prediction

Similarly to hERG and Nav1.5, we conducted an evaluation of the most effective predictive model for Cav1.2 cardiotoxicity, using various combinations of molecular features, on two external test sets. One notable observation, distinguishing Cav1.2 from other targets, is the significant disparity in predictive performance between the Cav-70 and Cav-60 test sets (as depicted in Table 5 and Figure S12). This discrepancy can be attributed to the size of the available data we could gather for training, as compared to Nav1.5 and hERG, the training dataset for Cav1.2 was considerably smaller.

**Table 5.** Cav1.2 toxicity prediction performance of each feature combination using the deep learning model on the two external test sets Cav-70 and Cav-60.

|  | Cav-70 |  |  |  |  |  | Cav-60 |  |  |  |  |  |
| --- | --- | --- | --- | --- | --- | --- | --- | --- | --- | --- | --- | --- |
| Features used | AC | F1 | SN | SP | CCR | MCC | AC | F1 | SN | SP | CCR | MCC |
| Fingerprints | 82.7 | 87.5 | 94.2 | 62.1 | 78.1 | 61.6 | 59.7 | 70.6 | **73.2** | 33.3 | 53.3 | 6.8 |
| Descriptors | 85.2 | 89.1 | 94.2 | 69.0 | 81.6 | 67.2 | 64.5 | 71.8 | 68.3 | 57.1 | 62.7 | 24.5 |
| Graph | 75.3 | 81.5 | 84.6 | 58.6 | 71.6 | 44.9 | 59.7 | 66.7 | 61.0 | 57.1 | 59.1 | 17.2 |
| Fingerprints + Descriptors | **86.4** | **90.1** | 96.2 | 69.0 | 82.6 | **70.2** | **69.4** | **75.9** | **73.2** | **61.9** | **67.5** | **34.1** |
| Fingerprints + Graph | 84.0 | 88.7 | **98.1** | 58.6 | 78.3 | 65.4 | 59.7 | 69.1 | 68.3 | 42.9 | 55.6 | 11.0 |
| Descriptors + Graph | 80.2 | 85.2 | 88.5 | 65.5 | 77.0 | 56.0 | 62.9 | 70.9 | 68.3 | 52.4 | 60.3 | 20.1 |
| All Combined | **86.4** | 89.9 | 94.2 | **72.4** | **83.3** | 70.0 | 64.5 | 71.8 | 68.3 | 57.1 | 62.7 | 24.5 |

Notwithstanding the data limitations, the models exhibited impressive results for the Cav-70 test set across all feature combinations, achieving an average accuracy of approximately 82% and an F1-score above 85%. This suggests that the task of predicting Cav1.2 cardiotoxicity is relatively easy compared to the previous two targets. When considering individual feature sets and all evaluation metrics, we can rank the best-performing features from worst to best as follows: graph representation is the weakest, followed by fingerprints, and then molecular descriptor representations. Descriptors demonstrate high informative information in this task compared to the other two features. When combined with fingerprints, the performance is significantly enhanced. Similar to Nav1.5, the optimal model for Cav1.2 combines both fingerprint and descriptor-based representations. We have named this model CToxPred-Cav1.2, with an average standard error of ±3.8%. For the confusion matrices detailing the predictions of CToxPred-Cav1.2 on the two external test sets, please refer to Supplementary Figure S13.

## 4. Conclusion

Cardiac voltage-gated ion channels are collectively responsible for generating the action potential, which is required for cardiac cells’ contraction. Notably, the hERG, Nav1.5, and Cav1.2 ion channels are key components of the cardiac action potential, and their inhibition by drugs can cause severe cardiovascular complications. Therefore, accurate prediction of the potential cardiotoxic liability of these channels in drug interactions is crucial. This research addresses this need by assembling a large and comprehensive dataset of small molecules specifically tailored for this purpose. An important element of our effort is that the cardiotoxicity database is freely available as open access for further community development of machine cardiotoxicity prediction models. This is in contrast to previous efforts that have employed proprietary and private datasets that are not publicly accessible for the scientific community, such as GOSTAR [40].

Standard datasets are necessary for the proper comparison of different tools. We seek through this study to emphasize the importance of establishing a robust and comprehensive framework consisting of extensive and publicly available development and external test sets. Such a framework enables the scientific community to focus on developing AI models while facilitating the straightforward benchmarking of model performance; Hence avoiding over-optimistic results as presented earlier in the introduction in Konda et al. [26] and Doddareddy et al. [28] data as well as in the data leakage by CardioTox [21] observed in the sub-section 3.2.1 of the results section. Furthermore, in order to assess the generalizability of developed tools, it is necessary to evaluate on structurally dissimilar datasets. The more different the test data is from the training data, the lower the performance, as evidenced by the drop in performance from the Eval-70 to Eval-60 test sets for each of the targets.

Additionally, a deep learning model has been developed that uses multiple types of feature representations, including fingerprints, descriptors, and molecular graph representations. A thorough benchmarking of these representations has been conducted to identify the most effective combinations for each respective task. Overall, the results demonstrate that descriptors possess higher predictive power, resulting in improved models and enhanced generalizability, particularly for Nav1.5 and Cav1.2. The results have confirmed a finding by Jiang et al. [73], who did empirically demonstrate that on average the descriptor-based models outperform the graph-based models in terms of prediction accuracy and computational efficiency. In the case of hERG, fingerprints alone prove sufficient for discriminating blockers from non-blockers. However, for Nav1.5 and Cav1.2, the combination of fingerprints and descriptors yields the best performance. These results demonstrate how simple features can perform better than more complex ones, such as GNN-based features. As a result, we advocate for carefully evaluating various feature representations instead of immediately using the most complex deep learning models possible.

The database utilized in this study is intended to serve as a comprehensive framework for researchers in the field, enabling them to build predictive models and easily benchmark their results using consistent test sets. It is publicly available as open access on Zenodo at https://zenodo.org/record/8245086. As a result of this research, robust models for predicting cardiotoxicity for each ion channel have been developed and consolidated into a comprehensive tool named CToxPred.

In terms of future work, the collected dataset could be leveraged to develop regression models capable of directly predicting the estimated potency of molecular compounds. Additionally, structural modeling could be employed to validate the results. Other potential projects may explore the application of data augmentation techniques, particularly for Cav1.2 and Nav1.5, to assess improvements in predictions.

## Acknowledgement

The authors would like to thank Chanuka Fernando for his participation during the data collection.

## Notes

https://zenodo.org/record/8245086

https://github.com/issararab/CToxPred

### Competing Interest Statement

The authors have declared no competing interest.

