## Supplementary Online Material for "Benchmarking of Small Molecule Feature Representations for hERG, Nav1.5, and Cav1.2 Cardiotoxicity Prediction"

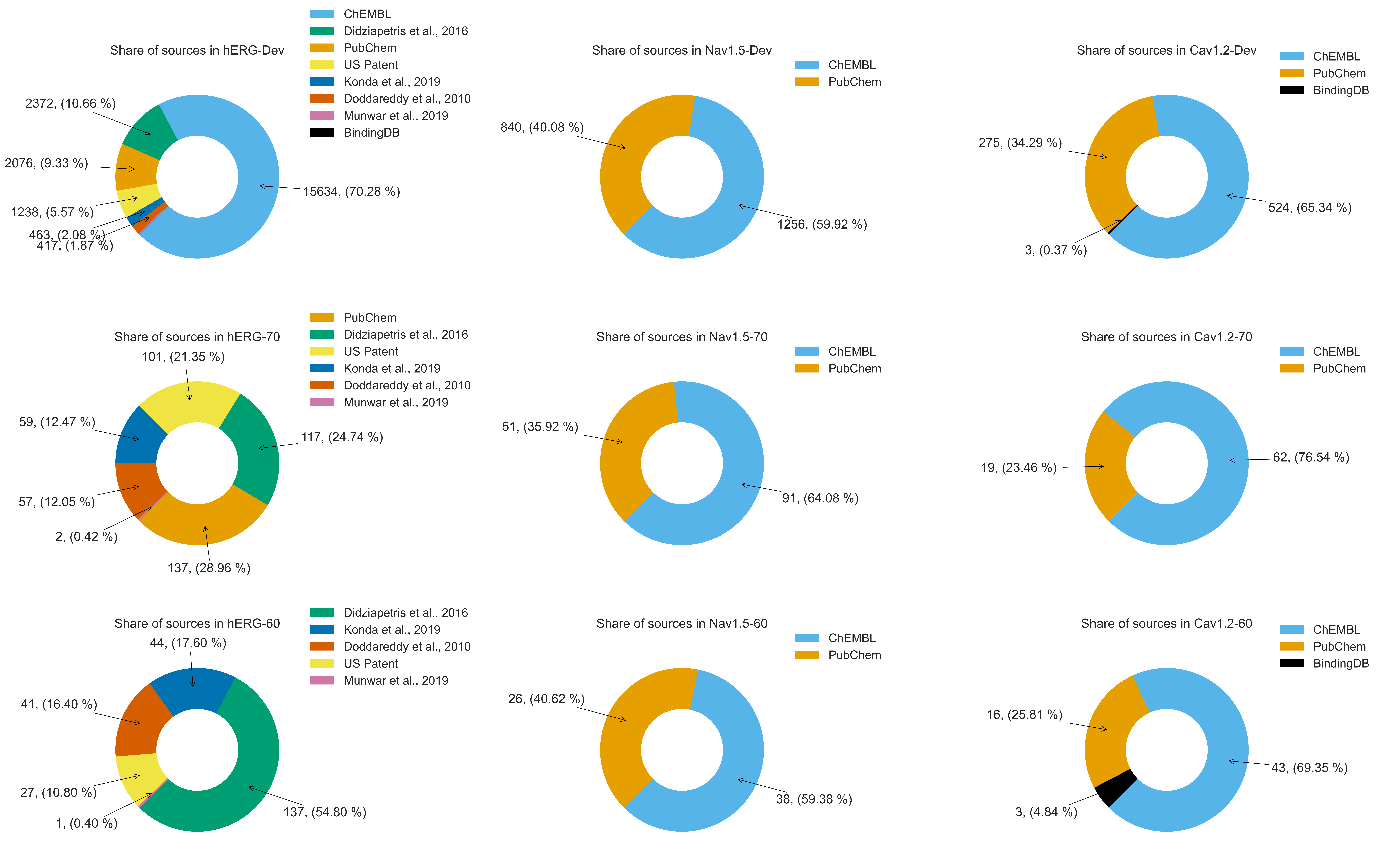

**Figure S1:** The composition of each set as derived from the upstream data sources in the final extensive cardiotoxicity database

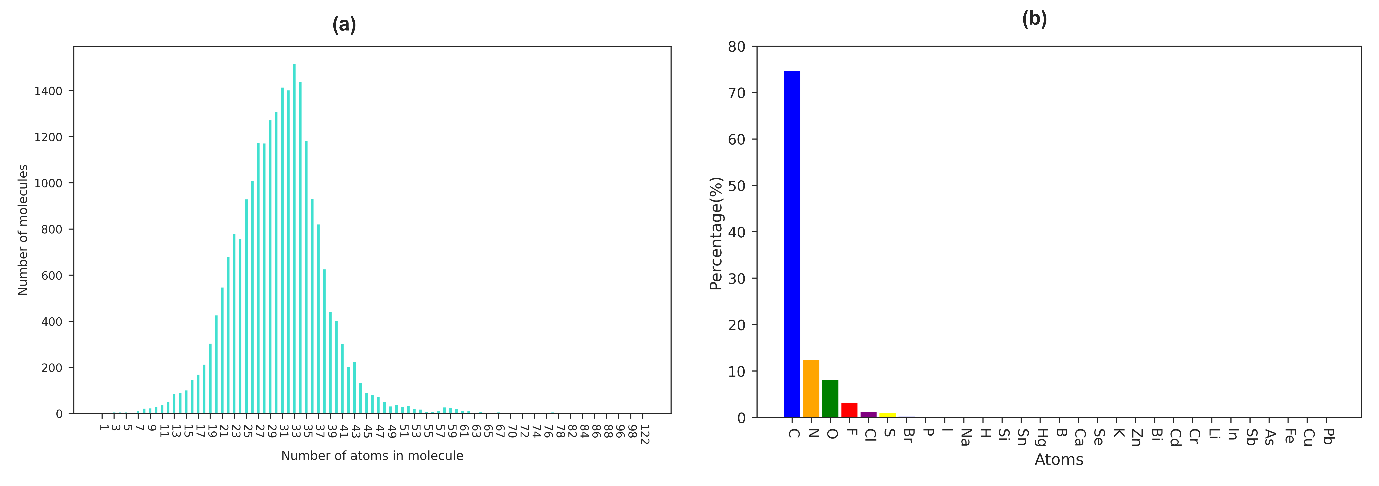

**Figure S2:** Atom composition analysis of molecules in our hERG development set. (a) represents a unimodal distribution of the total number of atoms in each molecule with a mean of around 30, while (b) showcases the atom composition distribution of the dataset

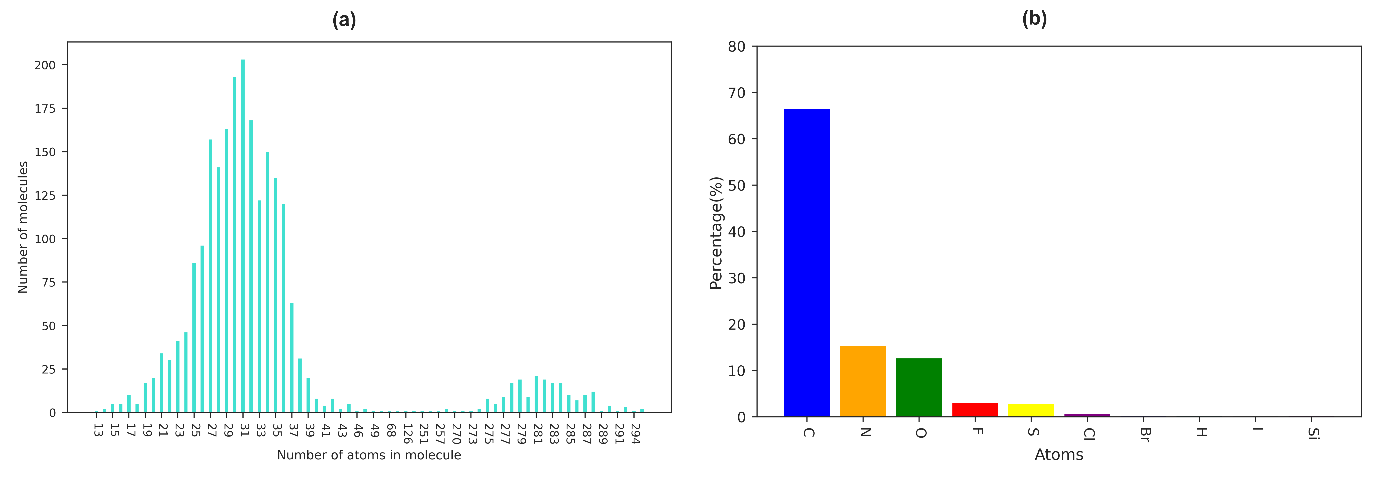

**Figure S3:** Atom composition analysis of molecules in our Nav1.5 development set. (a) represents a multimodal distribution of the total number of atoms in each molecule with an arithmetic mean around 52, while (b) showcases the atom composition distribution of the dataset

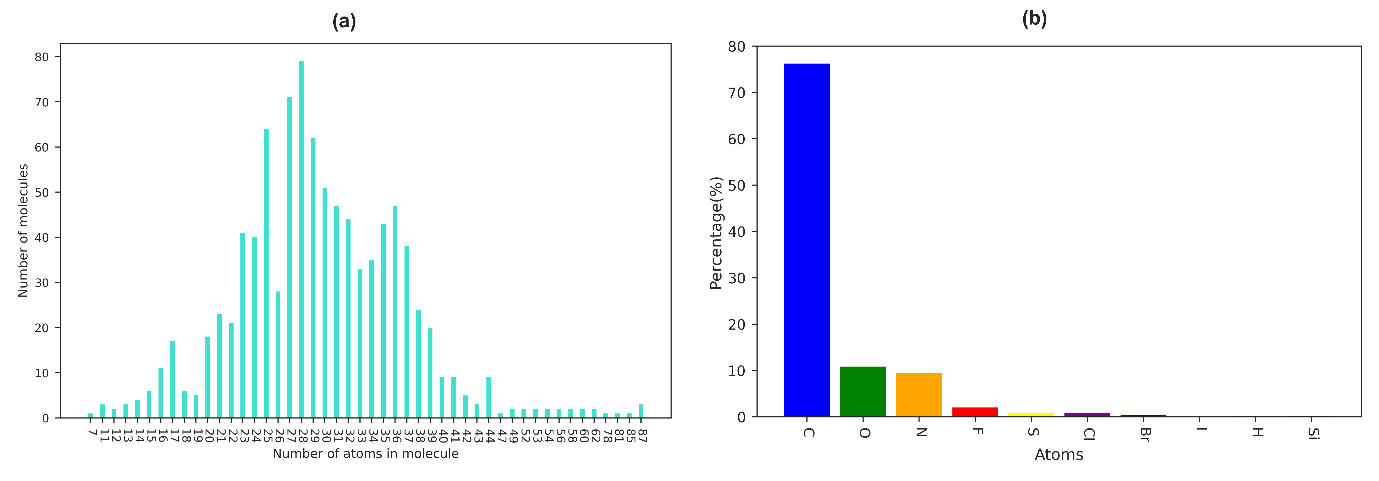

**Figure S4:** Atom composition analysis of molecules in our Cav1.2 development set. (a) represents a unimodal distribution of the total number of atoms in each molecule with a mean of around 30, while (b) showcases the atom composition distribution of the dataset

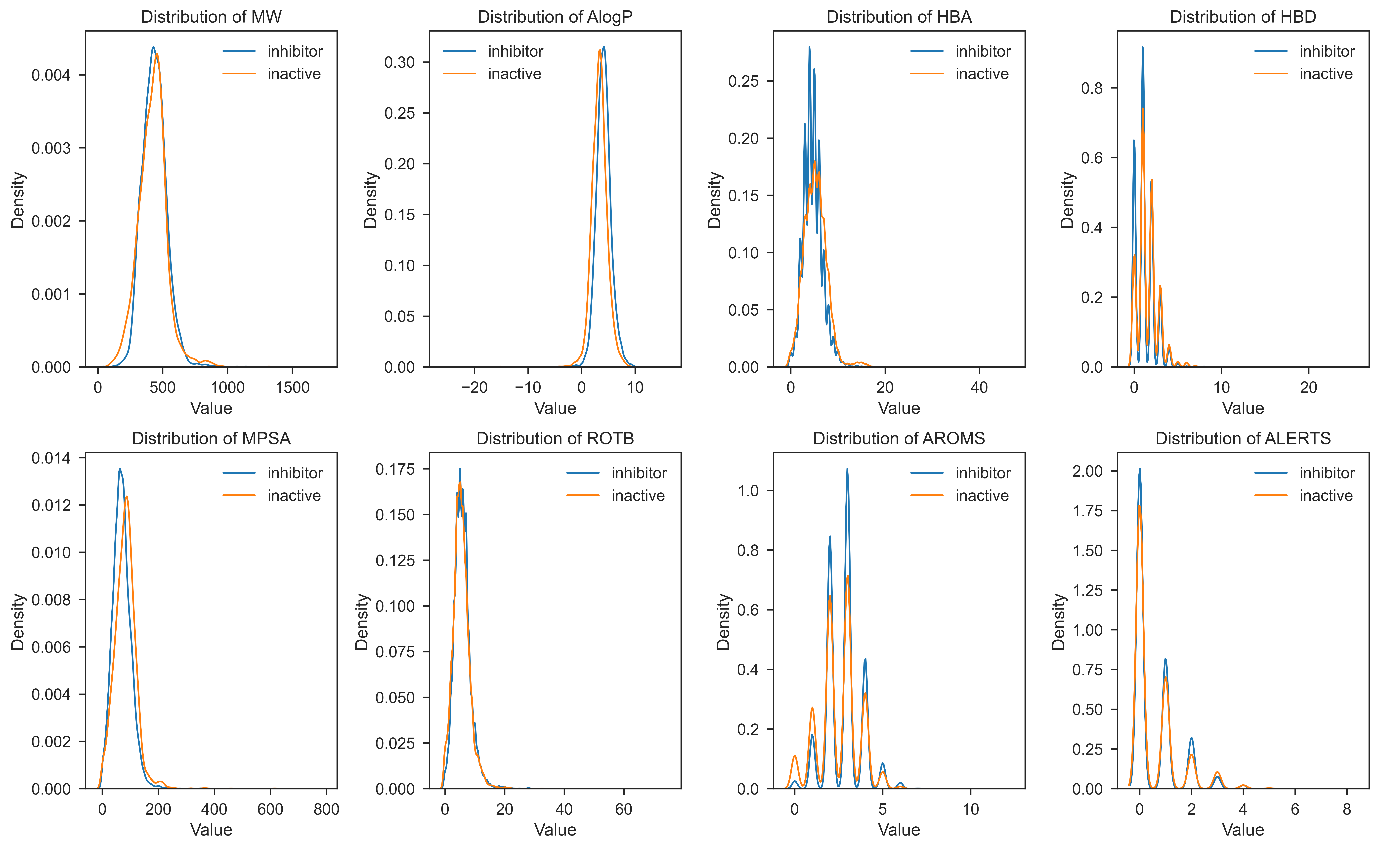

**Figure S5:** Distributions of the 8 physicochemical properties between inhibitor(blocker) and inactive(non-blocker) compounds in the hERG dataset

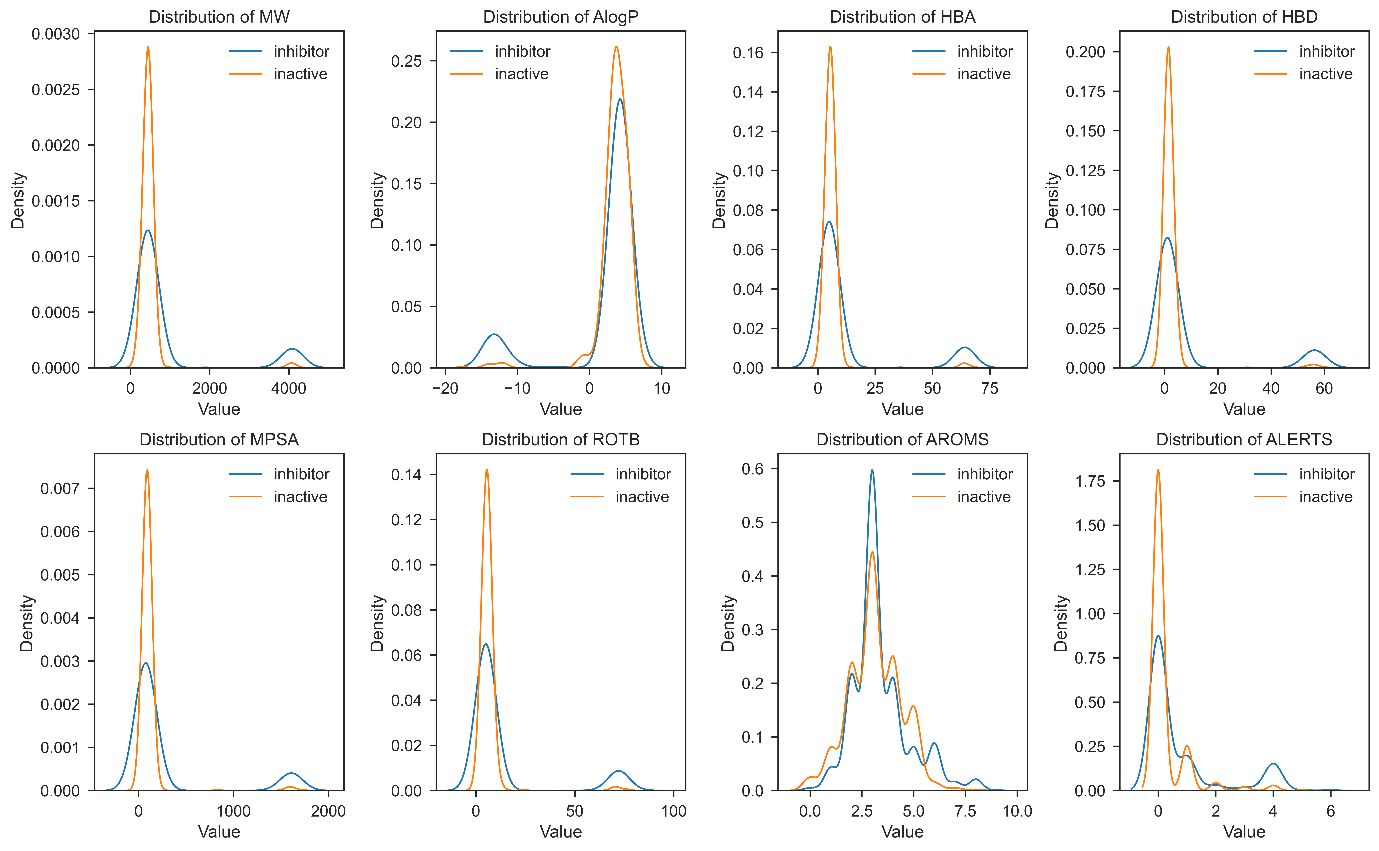

**Figure S6:** Distributions of the 8 physicochemical properties between inhibitor(blocker) and inactive(non-blocker) compounds in the Nav1.5 dataset

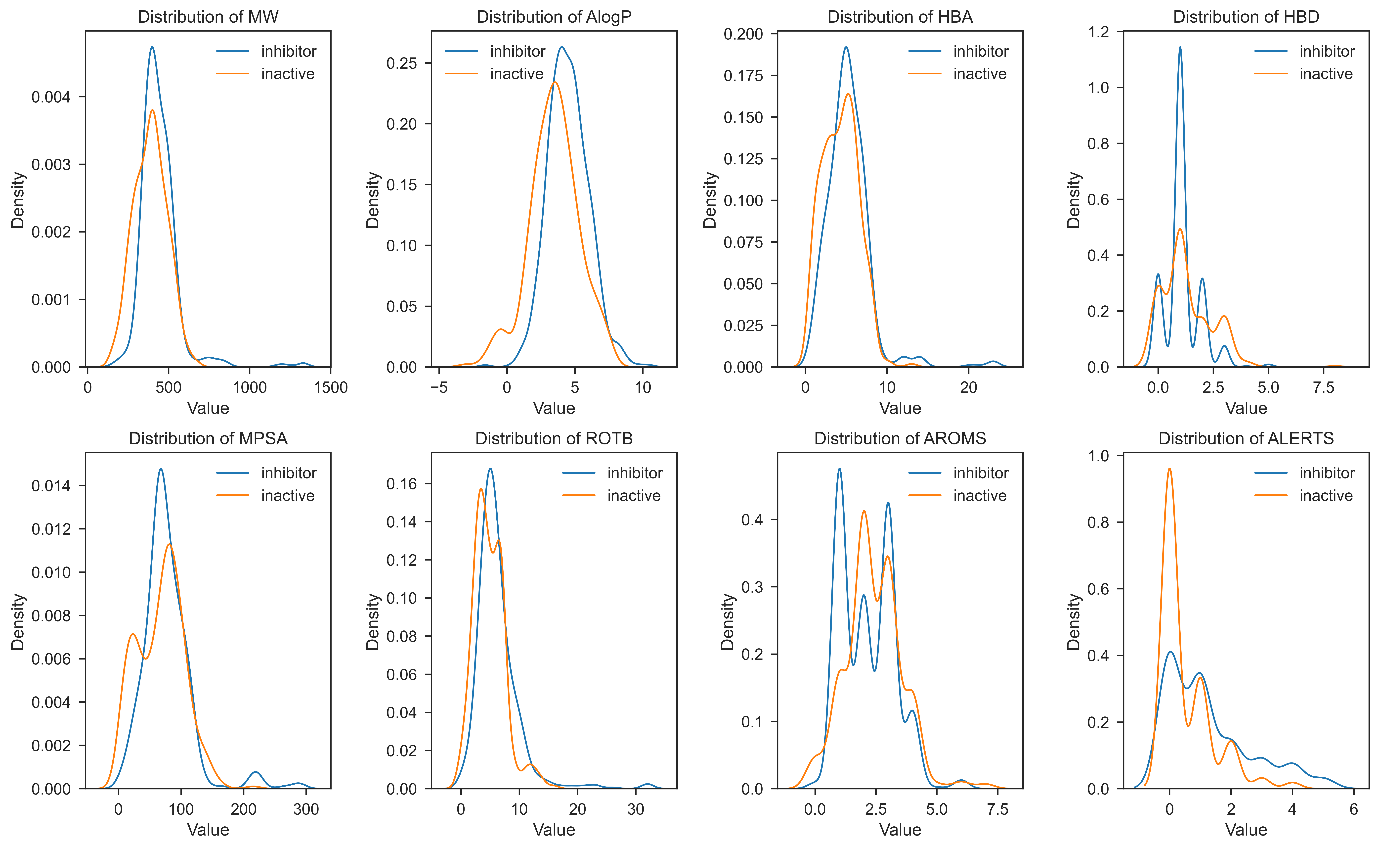

**Figure S7:** Distributions of the 8 physicochemical properties between inhibitor(blocker) and inactive(non-blocker) compounds in the Cav1.2 dataset

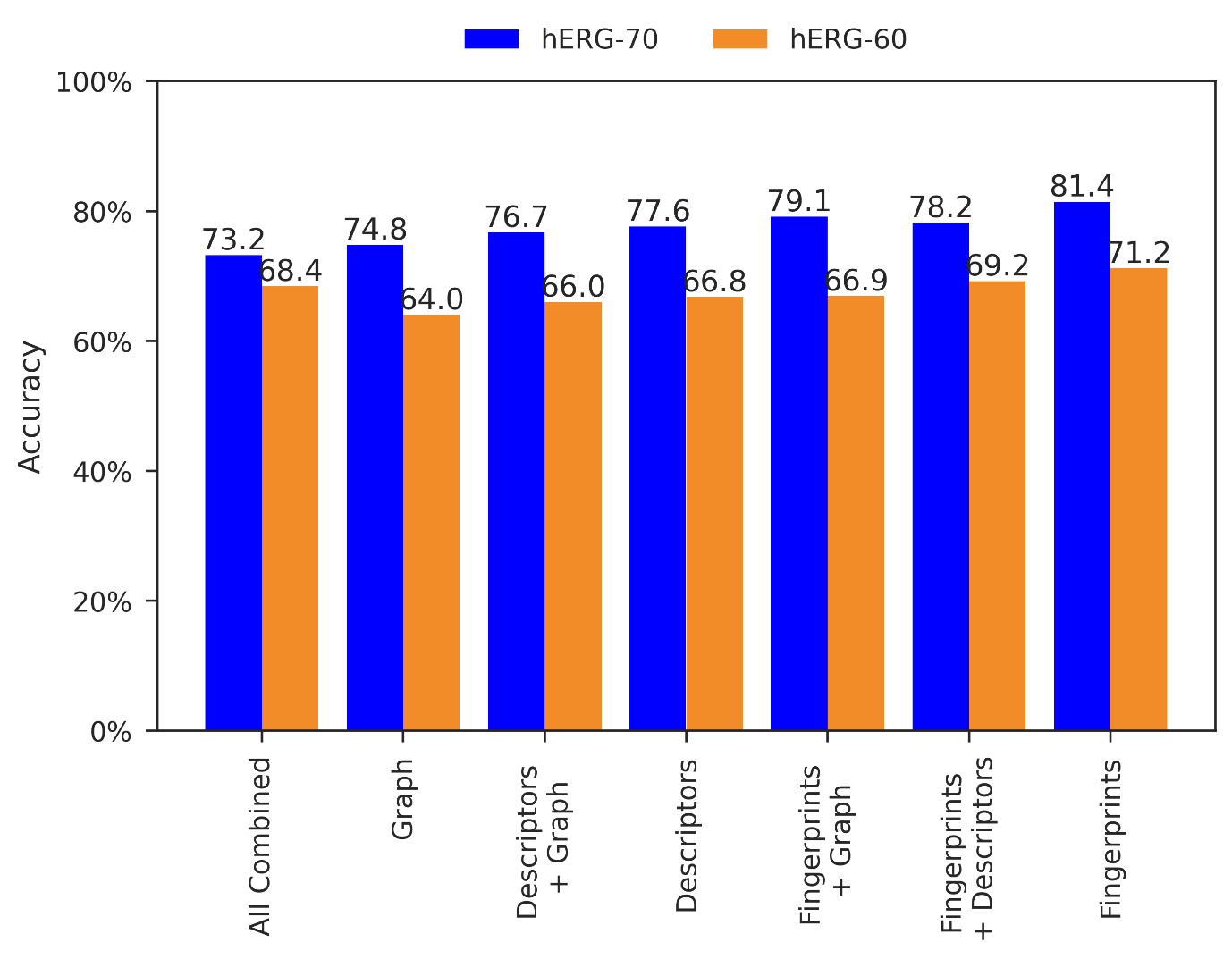

**Figure S8:** Accuracy performance comparison of different feature combinations on hERG liability prediction

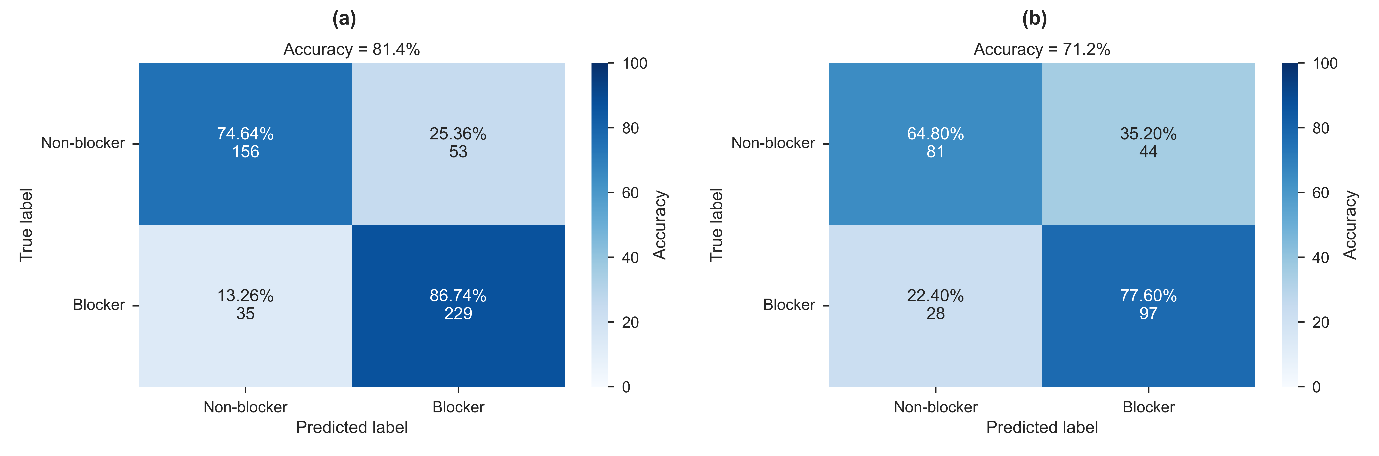

**Figure S9:** CToxPred-hERG (a) confusion matrix on the hERG-70 (b) confusion matrix on the hERG-60

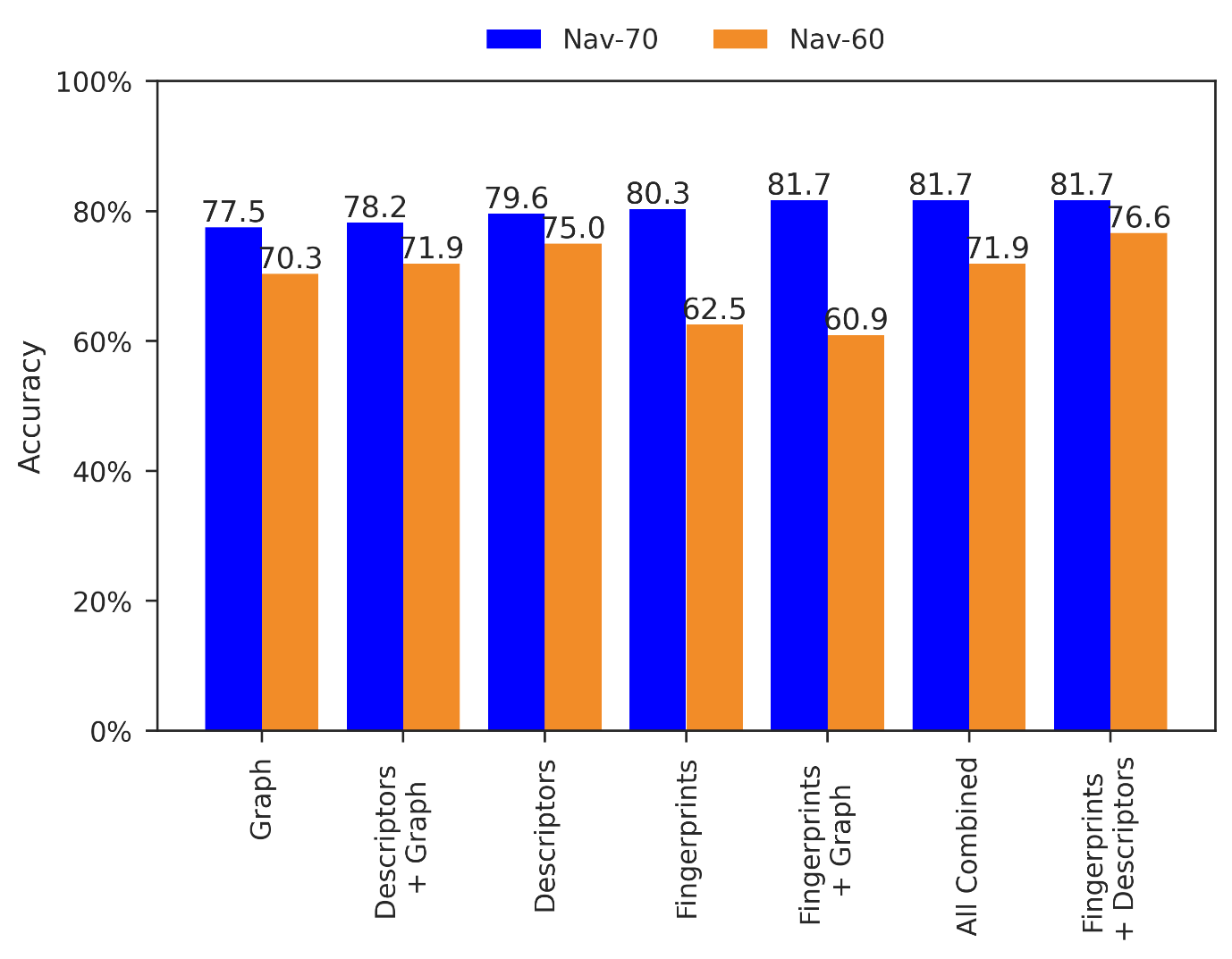

**Figure S10:** Accuracy performance comparison of different feature combinations on Nav1.5 liability prediction

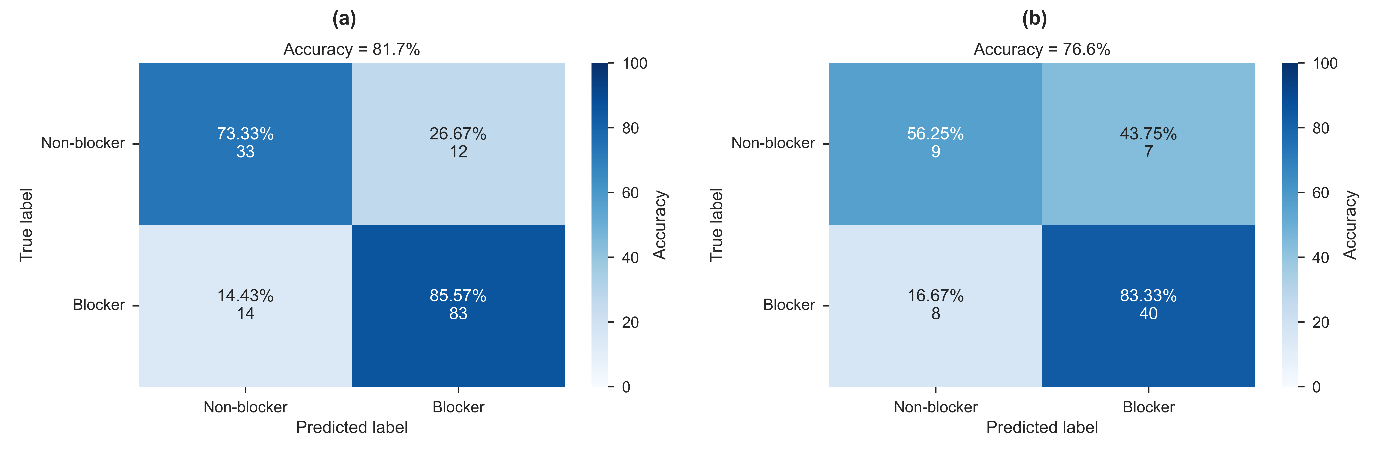

**Figure S11:** CToxPred-Nav1.5 (a) confusion matrix on the Nav-70 (b) confusion matrix on the Nav-60

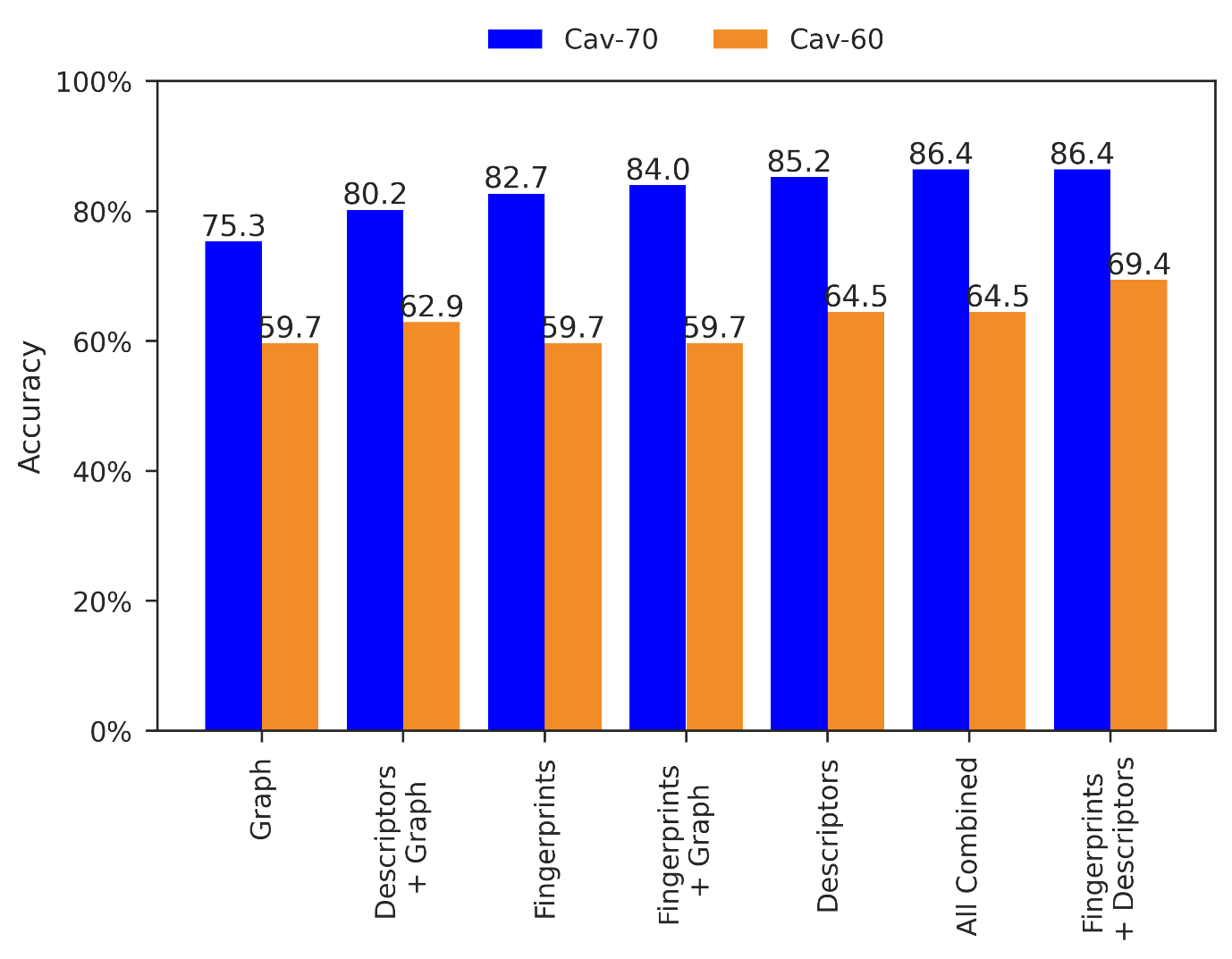

**Figure S12:** Accuracy performance comparison of different feature combinations on Cav1.2 liability prediction

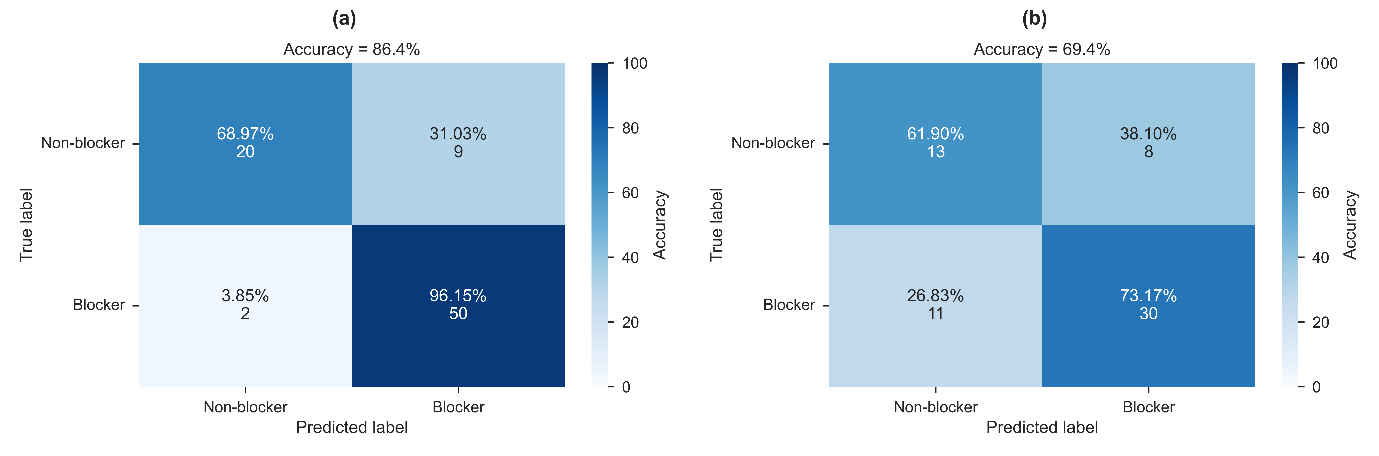

**Figure S13:** CToxPred-Cav1.2 (a) confusion matrix on the Cav-70 (b) confusion matrix on the Cav-60
